# Parentage assignment with genotyping-by-sequencing data

**DOI:** 10.1101/270561

**Authors:** Andrew Whalen, Gregor Gorjanc, John M Hickey

## Abstract

In this paper we evaluate using genotype-by-sequencing (GBS) data to perform parentage assignment in lieu of traditional array data. The use of GBS data raises two issues: First, for low-coverage GBS data, it may not be possible to call the genotype at many loci, a critical first step for detecting opposing homozygous markers. Second, the amount of sequencing coverage may vary across individuals, making it challenging to directly compare the likelihood scores between putative parents. To address these issues we extend the probabilistic framework of Huisman (2017) and evaluate putative parents by comparing their (potentially noisy) genotypes to a series of proposal distributions. These distributions describe the expected genotype probabilities for the relatives of an individual. We assign putative parents as a parent if they are classified as a parent (as opposed to e.g., an unrelated individual), and if the assignment score passes a threshold. We evaluated this method on simulated data and found that (1) high-coverage GBS data performs similarly to array data and requires only a small number of markers to correctly assign parents and (2) low-coverage GBS data (as low as 0.1x) can also be used, provided that it is obtained across a large number of markers. When analysing the low-coverage GBS data, we also found a high number of false positives if the true parent is not contained within the list of candidate parents, but that this false positive rate can be greatly reduced by hand tuning the assignment threshold. We provide this parentage assignment method as a standalone program called AlphaAssign.

## Introduction

In this paper we evaluate the performance of using genotype-by-sequence (GBS) data to perform parentage assignment in commercial plant and animal breeding settings. Having accurate parentage information is important for many routine breeding applications, such as reducing the cost of genotyping through pedigree-based imputation (Huang et al., 2012), reducing the bias of genomic estimates of breeding values (Solberg et al., 2009), and combining genotyped and non-genotyped individuals into a joint analysis (Legarra et al., 2009). When the parents of an individual are not recorded, parentage assignment algorithms can use genetic data to reconstruct parent-child relationships. Much of the previous work on parentage assignment has focused on the case where the genetic data was generated from microsatellite markers or more recently from SNP arrays (Rohrer et al., 2007; Fisher et al., 2009; Riester et al., 2009; Tokarska et al., 2009). In the case of SNP arrays between 50 and 700 markers are required to accurately assign parents and rule out false assignments (Rohrer et al., 2007; Strucken et al., 2016; Fisher et al., 2009; Tortereau et al., 2017). GBS is a flexible alternative to arrays, particularly for species that may not have a well-established reference genome, or where a suitable array has not been developed. However, the performance of using GBS data for parentage assignment – to our knowledge – is not well understood.

The primary challenge for using GBS data is the potentially high uncertainty in the true genotype of an individual based on the observed genetic data. In a GBS platform, a restriction enzyme is used to cut DNA into fragments that are then sequenced (Baird et al., 2008; Davey et al., 2011; Elshire et al., 2011). This means that unlike arrays, which produce called genotypes, GBS produces read counts for the reference and alternative alleles. For high-coverage GBS data the underlying genotype can easily be called from the read counts. For low-coverage GBS data calling genotypes is more difficult, particularly on loci which only receive a few reads. Distinguishing between heterozygous and homozygous loci is particularly challenging. If GBS produces two reads for the reference alleles and zero reads for the alternative allele, this could indicate that the individual is homozygous for the reference allele, or the individual could be heterozygous and their reference allele was sequenced twice. The difficulty in calling homozygous loci makes parentage assignment particularly difficult because many parentage assignment algorithms, either explicitly or implicitly, rely on finding opposing homozygous loci to filter out putative parents. In addition, the lack of opposing homozygous loci may increase false positive rate of parentage assignment if the true parent is not in the list of putative parents, since full sibs or half sibs of the true parent may appear to be more related to the individual than expected by chance (Meagher and Thompson, 1986).

Likelihood based methods (e.g., Kalinowski et al., 2007; Riester et al., 2009) are one solution to handle genetic data with high uncertainty. In a likelihood based method, parentage assignment is based on the likelihood of an individual’s genotype conditioned on the putative parent’s genotype. If the genotypes of either the individual or the putative parent cannot be assessed accurately, this likelihood score can be calculated by marginalizing over possible genotypes. Likelihood methods work well in cases where all individuals have the same amount of genetic data (e.g., same number of markers or sequencing coverage), but may break down when individuals are genotyped at a different number of markers or at different coverage levels. An example of this could be two putative parents with array data. Suppose the first putative parent was genotyped at 50 markers that overlap with the child, and the second was genotyped at 1,000 markers that overlap with the child. If both parents were heterozygous at all loci and we assume that the loci are not linked, then the likelihood value for the first parent would be .5^50^ (each allele having a 50% chance of being transmitted), whereas the likelihood value for the second parent would be .5^1000^. These likelihood values are hard to compare against each other because they are calculated on different sets of markers. This problem can be solved by selecting a subset of markers that are genotyped in all putative parents (which may drastically reduce the amount of information available), or using the population allele frequency for the genotype at missing markers (which disadvantages individuals with missing values).

A third option, that may be more appealing for GBS data, is to instead change the parentage assignment problem into a relationship classification problem. With this framing, the goal of the algorithm is to classify the relationship between each putative parent and the focal individual (e.g., parent, grandparent, sibling, child). A putative parent is then assigned as the parent, if they are classified as a parent, pass an assignment threshold, and are the highest scoring parent out of the list of putative parents (Huisman, 2017; Riester et al., 2009). One of the main advantages of this approach is that the classification task (which is able to filter out most putative parents) only relies on the genetic information available for an individual and a putative parent and does not require direct comparison to other putative parents. This property is particularly appealing for GBS data where the amount of information on each individual may differ depending on the genotyping resources spent and the allele frequency of the loci with sequence reads.

In this paper we extend the parentage assignment method of Huisman (2017) to explicitly handle GBS data. We then evaluated its performance in a simulated animal breeding population. We found that, similar to array data, it is possible to obtain accurate parent assignment with a fairly small number of sequence reads (e.g., 0.1x coverage), but that ruling out false positives is harder, and that a sizeable number of false positives could occur for medium coverage (0.5-2x) GBS data on a large number of linked markers.

## Materials and Methods

Here we describe our approach for parentage assignment with GBS data. This work builds closely on the probabilistic framework of Huisman (2017), but we present the full model for completeness. To assign parents we first construct a series of proposal distributions for each putative parent based on the genotypes of a focal individual and it’s known relatives. These proposal distributions describe the expected genotypes for a relative as a function of their relationship with the focal individual (e.g., parent, full sib of the parent, unrelated). We then classify each putative parent into one of these relationships, and if it is classified as a parent, and the assignment score passes a threshold, we assign it as the parent. If there are multiple possible parents, the highest scoring individual is assigned. Although this algorithm was originally designed in the context of animals, it also works for diploid and allopolyploid plants.

To simplify the language, we assume that we are attempting to assign the father of a focal individual. For a given focal individual *i* and its mother *m* we calculate the probability that the putative parent *f* is the true father by:

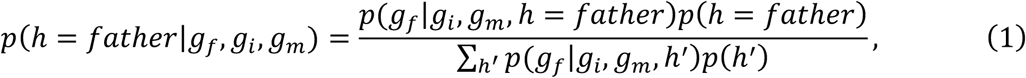

where *g_x_* is the genotype of individual *x*, *h* is the relationship between the focal individual *i* and the putative parent *f*, and the denominator is enumerated over the set of possible relationships *h*′. In the case where the genotypes of the mother are unknown we assume that her genotype probabilities are derived from Hardy-Weinberg Equilibrium.

In this paper we consider four possible relationships: that the putative parent is the true father, a full sib of the true father, a half sib of the true father, or unrelated. The conditional probability distributions for alternative relationships can be constructed via the generative framework we provide below. To simplify calculations, we assume that *p(h’)* is uniform over all possible relationships. In addition, we assume all markers segregate independently allowing *p*(*g_f_*∣*g_i_*, *g_m_*, *h*) to be calculated as the product of the probability of the putative parent’s genotype at each marker *k*:

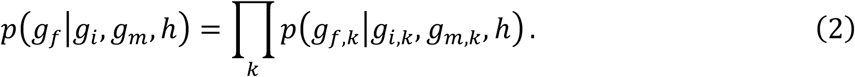

In the case of array data, and particularly GBS data, our assessment of the true genotypes, *g_f_*, *g_i_*, and *g_m_* may be noisy. To account for this noise we marginalize across possible genotypes based on observed genetic data ***d*** = *(d_i_, d_f_ d_m_)*:

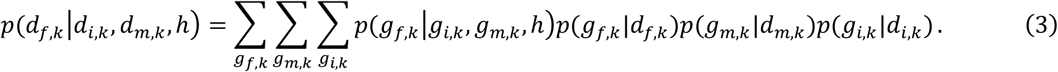

This model requires the calculation of two terms: (1) the genotype probabilities conditional on the observed data *p*(*g*_*x*,*k*_∣*d*_*x*,*k*_) and (2) the proposal distribution for an individual’s genotype based on their relationship with the focal individual *p*(*g*_*f*,*k*_∣*g*_*i*,*k*_,*g*_*m*,*k*_,*h*). We outline how to calculate both terms below.

### Evaluating genotype probabilities conditional on the observed data

In this model we assume that each marker is biallelic and has four possible phased genotypes, aa, aA, Aa, AA. With observed array data for marker *k, d_x,k_*, the conditional probabilities for each genotype *g*_*x*,*k*_ are:

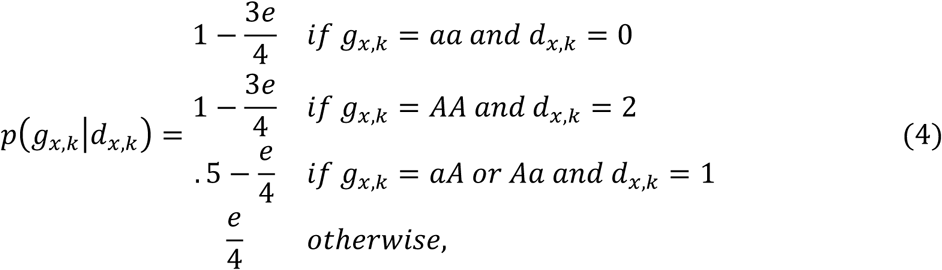

where *e* is the assumed genotyping error rate. This evaluation of individual genotype probabilities differs from Huisman (2017), where it is assumed that errors can only occur between homozygous and heterozygous states (and not between opposing homozygote states) and distinction is not made between two heterozygous genotypes. The genotype probabilities above correspond more closely to those commonly used in peeling (e.g., Whalen et al., 2017) and allow inferences to be made even when the genotyping error rate is high.

With observed GBS data for marker *k, d_x,k_*, the conditional genotype probabilities are:

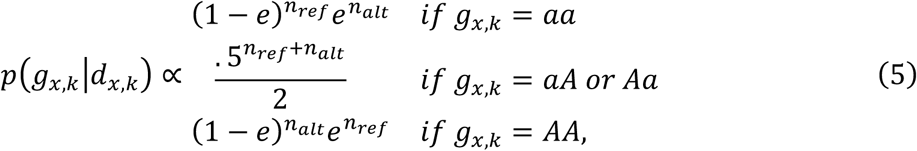

where *e* is the sequencing error rate, *n_ref_* is the number of sequence reads supporting the reference allele and *n_alt_* is the number of sequence reads supporting the alternative allele. The genotype probabilities in Equation 5 do not sum to one, and so the probabilities need to be normalized for each allele. Equation 4 is consistent with previous work on parentage assignment with array data (Kalinowski et al., 2007; Huisman, 2017), while Equation 5 is consistent with previous work on imputation with GBS-like data (Li et al., 2010; VanRaden et al., 2015; Whalen et al., 2017).

### Generating proposal distributions via single locus peeling

We generate proposal distributions *p*(*g*_*f*,*k*_∣*g*_*i*,*k*_,*g*_*m*,*k*_, *h*) for the genotype probabilities of each relationship via single locus peeling (Elston and Stewart, 1971). Single locus peeling provides a rich generative model for estimating the genotype probabilities of un-genotyped relatives based on the genotypes of an individual and a known parent. Although our presentation differs from Huisman (2017) it results in the same distributions. Under this framework, we calculate the genotype probabilities for three relatives: the father, a full-sib of the father, and a half-sib of the father. These probabilities are calculated by first estimating the genotype probabilities for the father, peeling up to the paternal grandparents, and finally peeling down to the full sib and the half sib of the father (Figure 1).

**Figure 1.**
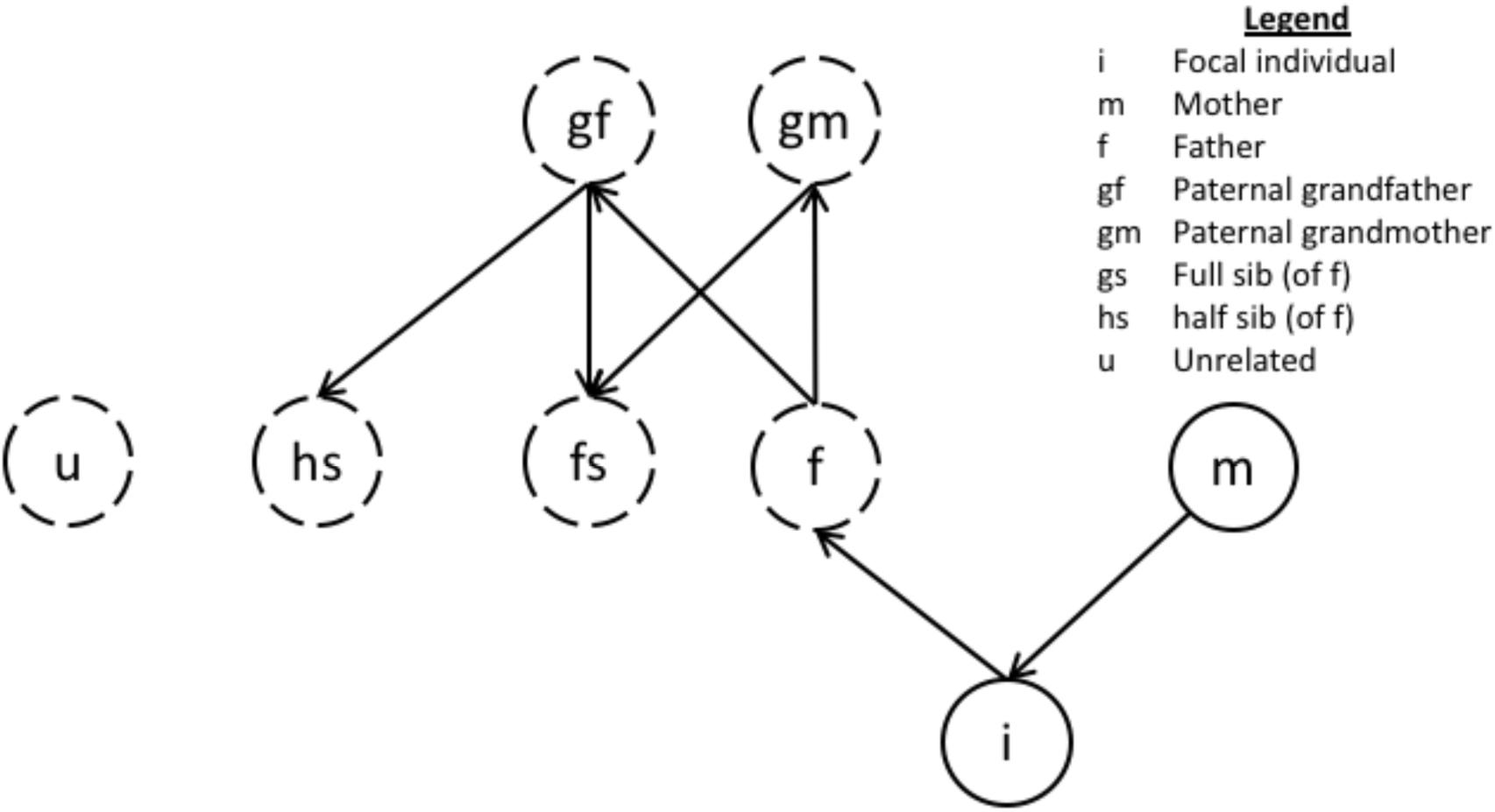
A graphical representation of the peeling order for the proposal distributions. The arrows represent the direction in which the peeling operations should be performed. Hardy-Weinberg equilibrium is used to generate the genotype distributions for the unrelated individual, the mother of the half sib, and if unknown, the mother’s genotype. Although this graphic assumes the mother is known and the father unknown, a symmetric picture could be constructed when the mother is unknown and father known.

Given genetic data on the focal individual *d_i_* and a mother *d_m_*, we can construct a proposal distribution for the father via:

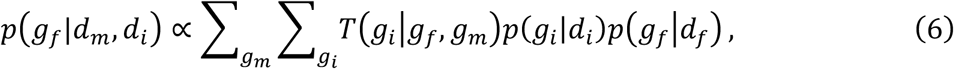

where *p*(*g_i_*∣*d_i_*) is given by Equation 4 or 5 above, and *T*(*g_i_*∣*g_f_*,*g_m_*) is the probability that the individual inherited genotype *g_i_* conditional on their parents having genotypes *g_f_* and *g_m_*, e.g., *T*(*g_i_* = *aA*∣*g_f_* = *aA*,*g_m_* = *AA*) = 0.5 (Marshall et al., 2003).

Using Equation 6, we can peel up to construct a joint distribution for the genotypes of the paternal grandparents (*gf*, *gm*):

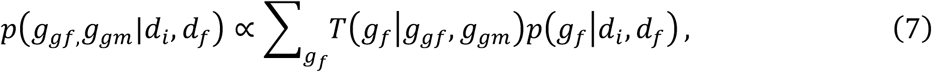

where *p*(*g_f_*∣*d_i_*, *d_f_*) is given in Equation 6, above. We can then peel down to generate the proposal distributions for a full sib and a half sib of the father. The proposal distributions differs in whether the full joint distribution of both grandparents is used (full sib,*fs*), or if only one of the grandparents is used and the other parent assumed to have genotypes based on Hardy Weinberg Equilibrium (half sib, *hs*):

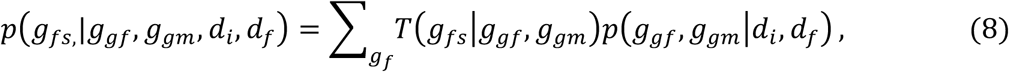

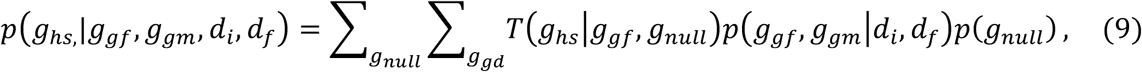

where *p*(*g_null_*) represents the probability of having a genotype if that genotype was drawn at random from the population.

The proposal distribution for an unrelated individual simply assumes that their genotypes are drawn at random from the population according to Hardy Weinberg Equilibrium:

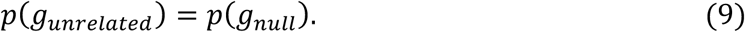

To assign a parent we calculated an assignment score for each putative parent:

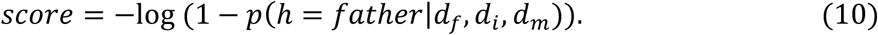

The score will be close to 0 if the individual is unlikely to be the father, and tends towards positive infinity with increasing evidence that the individual is the father. A putative parent was assigned as the true parent if its assignment score was the highest of the putative parents considered, and was higher than a pre-defined threshold.

Although the described process may seem computationally intensive, there are two features which simplify calculations. First, because the proposal distributions depend only on the focal individual and its known parent, the proposal distributions only need to be calculated once and can be re-used for all putative parents of the focal individual. Second, peeling can be performed efficiently as a series of tensor operations on the genotypes of focal individual and its known parent, filtered through the inheritance matrix *T*, which allows us to take advantage of linear algebra libraries.

### Simulated data

The simulated data modelled a livestock population. We initially sampled a set of genomes with 20 chromosomes using the Markovian coalescent simulator MaCS (Chen et al., 2009). For this we assumed that each chromosome is 10^8^ bp long, a per site mutation rate is 2.5 × 10^−8^, a per site recombination rate is 1.0 × 10^−8^, and that effective population size changed over time. Based on estimates for the Holstein cattle population (Villa-Angulo et al., 2009), we set the effective population size to 100 in the final generation of the coalescent simulation and to 1,256, 4,350, and 43,500 at respectively 1000, 10,000, and 100,000 generations ago, with linear changes in between. We then used the sampled chromosomes to initiate a population of 1,000 animals with equal sex proportions. We bred this population for 5 generations. In each generation, we selected 10 males and mated them at random to 100 females. Each potential focal individual therefore had 1 true father, 4 male full sibs of the father, and 45 male half sibs of the father. All individuals were genotyped at 50,000 markers. Subsets of these markers were used in different simulations as described below. Array data were simulated without any errors, due to the low error rate for modern SNP genotyping arrays (<1%; e.g., Kalinowski et al., 2007). In addition to array data, we generated low-coverage GBS data for the last two generations of individuals. We assumed that the GBS method targeted the same loci as the genotyping array and was performed at coverage levels between 0.1x to 10x. For each coverage level, the number of sequence reads at a given marker was generated via a Poisson distribution with mean equal to the coverage level. Each read randomly sampled one of the two alleles at a marker. The read sampling process also included a small sequencing error rate of 0.1%. We generated the simulated data using the R package AlphaSimR (Gaynor et al.), which is available at www.alphagenes.roslin.ed.ac.uk/AlphaSimR.

### Scenarios

We evaluated the accuracy of parent assignment for the last generation of 1,000 individuals across 4 different scenarios. In the first scenario (a) we analysed the accuracy of performing parent assignment when:

- the mother was known and genotyped,
- all of the male full- and half-sibs along with 50 other individuals (total of 100 potential parents) were putative parents,
- and either both the parents and progeny had array data, the parents had array data and the progeny had GBS data, or both the parents and the progeny had GBS data.

These sub-scenarios span a spectrum of possible practical settings. The sub-scenario where the parents had array data but the progeny had GBS data may represent either the case where the progeny are initially genotyped with a low-cost GBS platform and any selected parents are re-genotyped with an array, or it may represent the case where pedigree information was used to impute and accurately call parental genotypes. In the remaining scenarios we focused on the case where both parents and progeny had GBS data and analysed (b) the impact of knowing and genotyping the known alternative parent, (c) the impact of restricting the pool of putative parents to either 100 unrelated individuals, 45 half sibs, or the 4 full sibs, and (d) examined how the false positive rate changed depending on the threshold used for assignment (see below).

In each scenario we performed three evaluations. To evaluate the overall accuracy, we assumed the true parent was included in the list of putative parents, and evaluated accuracy by the number of times the top parent was the true parent. To evaluate the true positive rate we included the true parent in the list of putative parents, but assigned the top scoring parent only if it passed an assignment threshold. To evaluate the false positive rate, we excluded the true parent from the list of putative parents and counted the number of times the top scoring parent passed the assignment threshold. The first evaluation represented a case where we know the true parent is included in the list of putative parents (e.g., groups of females cohabitating with multiple males or artificial insemination using polyspermic matings). The second and third evaluations were designed to assess performance when we are not sure whether or not the true parent is included in the list of potential parents (e.g., natural service sires or wild populations).

### Software

Parentage assignment was performed using AlphaAssign (http://www.alphagenes.roslin.ed.ac.uk/alphasuite-softwares/alphaassign/) which, implements the described algorithm. AlphaAssign has three run-time parameters: (i) an assumed genotyping error rate for array data, (ii) an assumed sequencing error rate for GBS data, and (iii) an assignment threshold to determine the required score to assign a putative parent as a parent. Throughout this paper we assumed a 1% genotyping error rate, a 0.1% sequencing error rate, an assignment threshold of 10 (determined via pilot simulations) although we varied the assignment threshold in the last set of simulations.

## Results

### Parent assignment with array and GBS data

First we examined the number of markers required for accurate parentage assignment when both parents and progeny were genotyped with array data. If the true parent was included in the list of putative parents (and an assignment threshold was used), 100 markers were required to obtain 100% parentage assignment accuracy. If the true parent was excluded from the list of putative parents, the false positive rate was less than 0.1% if there were between 50 to 350 markers, and there were no false positives when there were more than 500 markers.

Unlike array data where the number of markers can be more easily varied, for GBS data the number of markers is usually determined by the choice of restriction enzymes while the amount of coverage obtained on each individual can be varied. Because of this we focused on the required coverage level to accurately assign parents based on a fixed number of markers. Figure 2 shows the accuracy and false positive rates based on the amount of coverage allocated to each progeny, stratified by the number of markers that this coverage is spread over. Because performance with array data was nearly identical to that with 10x GBS data we did not include array data in Figure 2.

**Figure 2.**
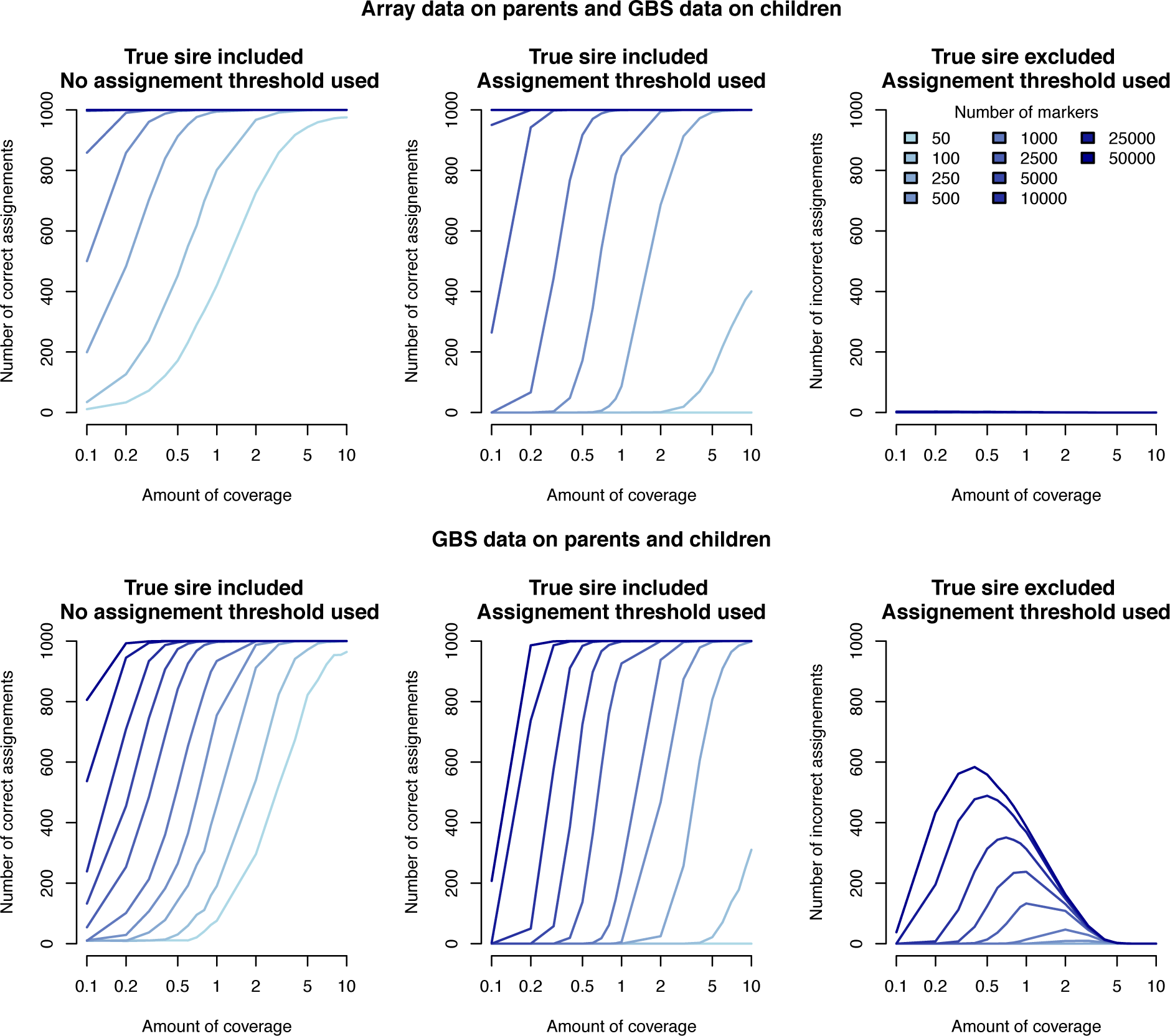
Parentage assignment performance when array or GBS data was available for the parents and GBS data was available for the progeny. The left panels give the number of correct assignments (for 1000 progeny) when the true parent was on the list of putative parents and no assignment threshold was used – the top scoring parent was assigned. The middle panels give the number of correct assignments when the true parent was on the list of putative parents and assignment threshold was used. The right panels give the number of incorrect assignments when the true parent was excluded from the list of putative parents.

We evaluated the performance of parentage assignment when the parents were genotyped with array data and the progeny were genotyped with GBS data. If the true parent was included in the list of putative parents, a coverage of 0.4x was required to obtain 100% accuracy when there were 50,000 GBS markers. The required coverage increased to 1x for 5,000 markers, and to 2x for 1,000 markers. If the true parent was excluded from the list of putative parents, we found that the false positive rate was less than 0.2% in all cases.

The accuracy of parentage assignment decreased when both the parents and progeny had GBS data. If the true parent was included in the list of putative parents, a coverage of 0.4x was required to obtain 100% accuracy when there were 50,000 GBS markers. The required coverage increased to 2x for 5,000 markers, and to 5x for 1,000 markers. If the true parent was excluded from the list of putative parents, we found that the false positive rate was as high as a 60%. These false positives were clustered on low to medium coverage GBS data (0.1 - 3x) with a large number of markers (>1000).

### False positive assignments by relationship

Figure 3 stratifies the false positive rate based on whether unrelated individuals, half-sibs of the true parent, or full-sibs of the true parent were included in the list of putative parents. In line with expectations we found a high false positive rate (as high as 60% in some conditions) when only the full-sibs of the true parent were included as putative parents. This decreased to at most 35% when only the half-sibs of the true parent were included and to under 20% when only unrelated individuals were included. As seen previously, most of the false positives were occurred when there were a large number of markers and low to medium coverage GBS data.

**Figure 3.**
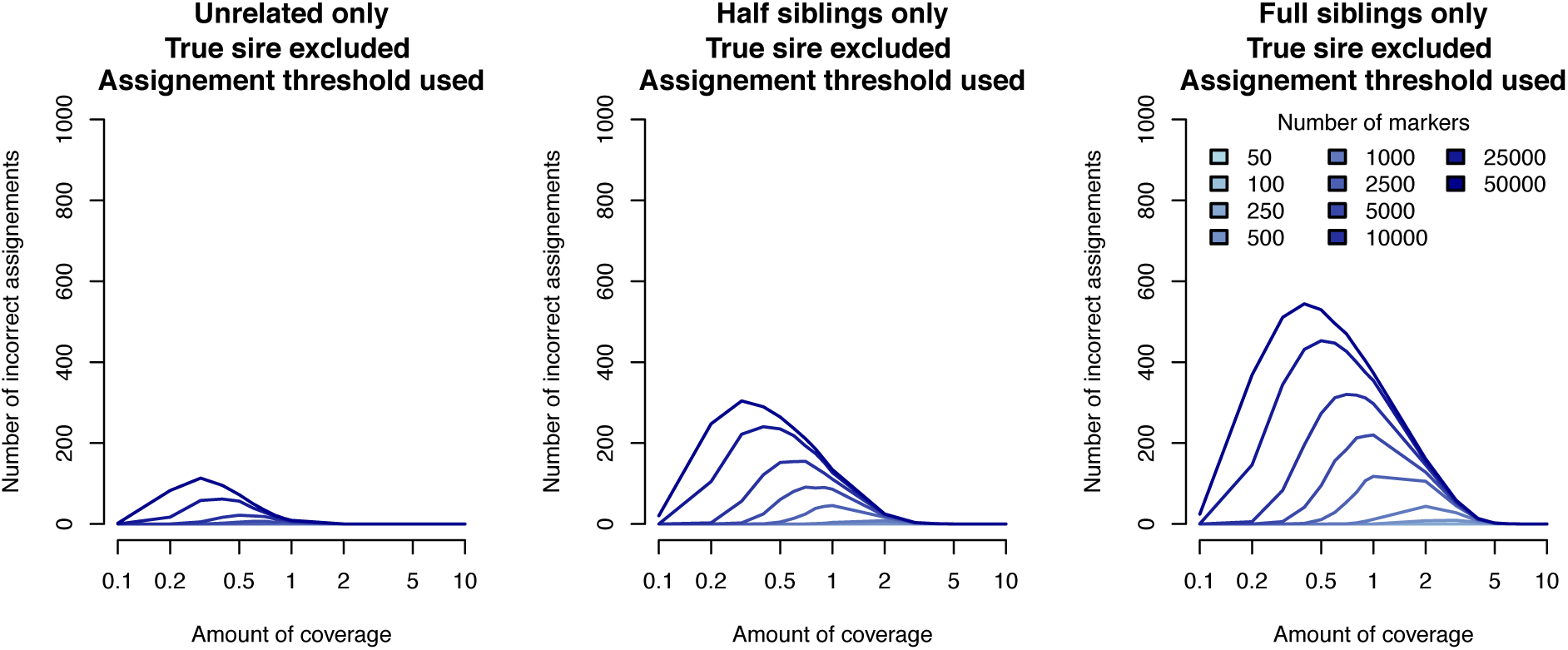
Number of false positive parentage assignments (for 1000 progeny) when GBS data was available for parents and progeny, the parent was excluded from the list of putative parents, assignment threshold was used, and the list of putative parents contained either 100 unrelated individuals (left panel), 45 half sibs of the true parent (middle panel), or 4 full sibs of the true parent (right panel).

### Parent assignment when neither parent is known

Figure 4 compares the performance of parentage assignment when one of the parents is known and genotyped compared to when neither parent is known or genotyped. We found that having one parent known and genotyped increased the accuracy of parentage assignment and decreased the number of false positives in all cases. The benefit was largest when both the progeny and parents had high-coverage GBS data.

**Figure 4.**
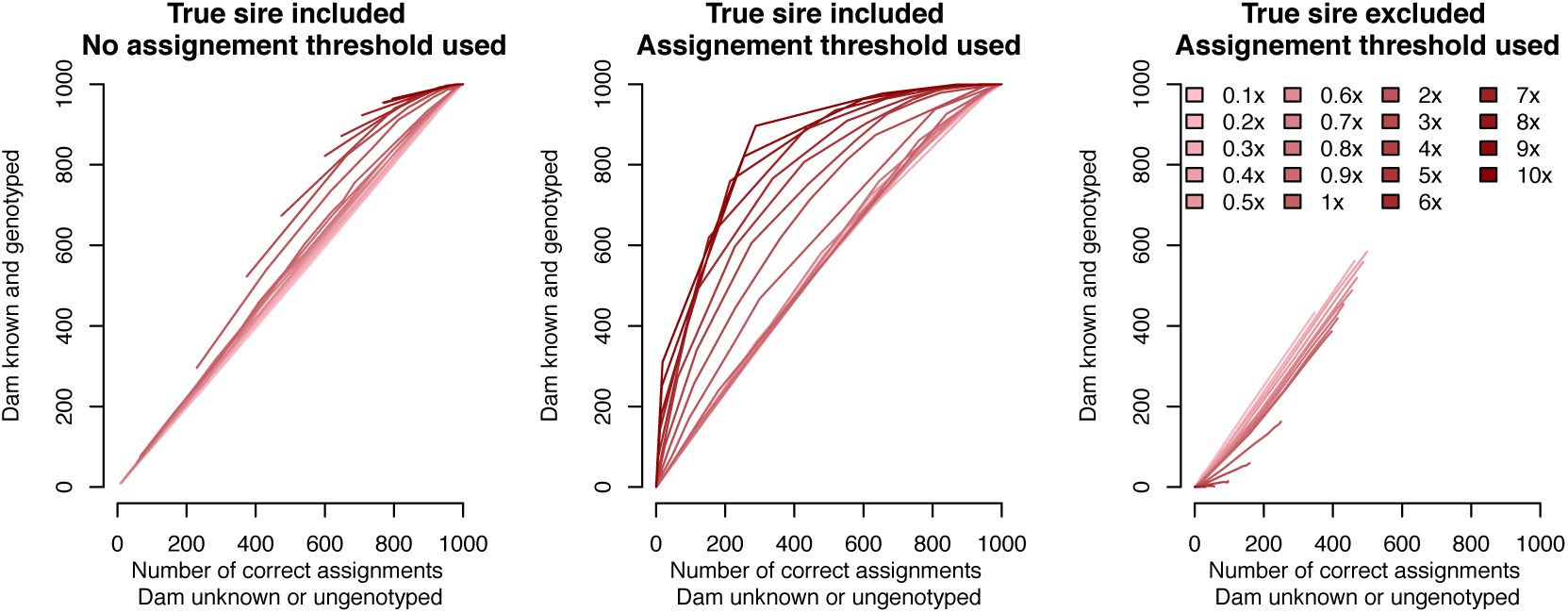
A comparison between the parentage assignment performance with one parent known and genotyped and no parent known at different GBS coverage levels (left and middle panes compare true positives while the right pane compares false positives).

### Controlling false assignments by modifying the threshold

Figure 5 shows the true positive rates and false positive rates for when sequencing resources were spread over 50,000 markers, as a function of the threshold used to assign a putative parent as the parent. We found that, compared to the results in Figures 2 and 3, it was possible to substantially reduce the false positive rate by increasing the assignment threshold, but that the ideal threshold depends on the total coverage. The relationship between the false positive and true positive rate is given as a receiver operating characteristic in Figure 5(c).

**Figure 5.**
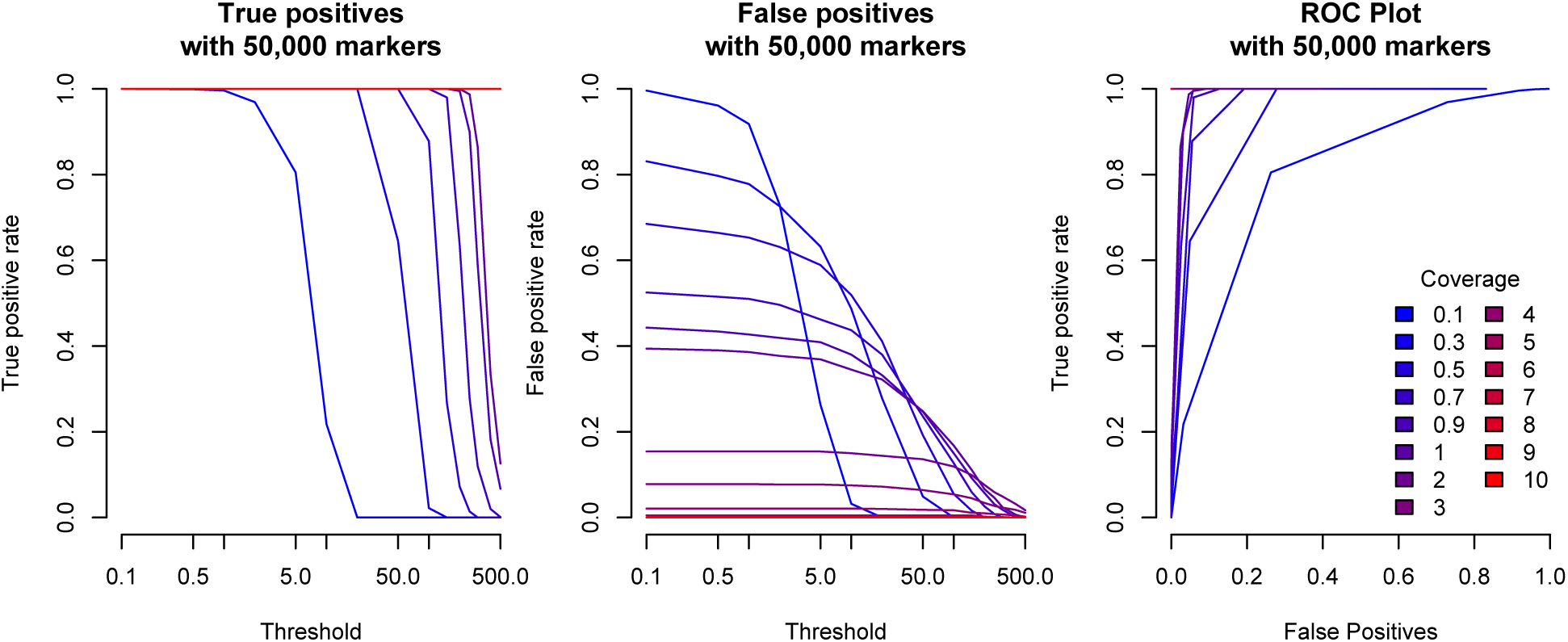
The rate of true positives, false positives, and the relationship between then when varying the total amount of coverage and the calling threshold.

### Timing

The algorithm took 3 minutes and 54 seconds to assign parents for 1000 progeny, each with 100 putative parents. The progeny and their parents were genotyped using GBS data across 5,000 markers. The algorithm scales linearly with the number of markers and the number of putative parents per individual.

## Discussion

In this paper we extended the parentage assignment method of Huisman (2017) to account for low-coverage sequence data and analysed the performance of parentage assignment when genotyping is performed via sequencing instead of the traditional genome-wide arrays. We found that high-coverage GBS data (i.e., 10x or higher) has the same performance as array data. We also found that low-coverage GBS data (as low as 0.1x) can be used to perform parentage assignment as long as it is obtained on a sufficiently large number of markers, but that there may be a large number of false assignments if the true parent is not included in the list of putative parents. The number of false positives could be reduced by modifying the threshold used to call assignments. In light of these results, we will discuss (1) the accuracy of parentage assignment, (2) potential extensions to control the false positive rate, and (3) the use of peeling to construct the proposal distributions in more detail.

### Parentage assignment accuracy with GBS data

A goal of this work was to quantify the amount of GBS data required to accurately perform parentage assignment. We found that, similar to array data, the total amount of data required is relatively low. For example, when using high-coverage GBS data between 100 to 200 markers are required to accurately assign parents. This is in line with previous estimates for array data (Rohrer et al., 2007; Strucken et al., 2016; Fisher et al., 2009; Tortereau et al., 2017), where between 50-700 markers were required. The differences in the exact number of markers required (100-200 compared to 50-700) is likely due to the structure of the underlying genetic data (i.e., number of chromosomes, minor allele frequency of the markers), and the assumption in this study that one of the parents was already known and genotyped.

In addition to being able to use high-coverage GBS data to perform parentage assignment, we found that low-coverage GBS data could also be used, provided it was spread across a larger number of markers. The increase in required number of markers is due to the lower information content at an individual loci for low-coverage GBS data, requiring the data to be pooled across a larger number of markers to achieve the same level of accuracy.

The results of this study suggest that GBS data – either high-coverage data on a small number of markers, or low-coverage data on a large number of markers – is an effective alternative to array data for performing parentage assignment. This result is particularly important given the emerging importance of GBS as an alternative for SNP array data, both in species where SNP arrays are available (e.g., De Donato et al., 2013; Brouard et al., 2017) and in those where SNP arrays have not been constructed (e.g., Robledo et al., 2017; Palaiokostas et al., 2018).

### Controlling the false positive rate

During our analysis of low-coverage data, we found an inflation of false positives when both the parents and the progeny had GBS data. These false positives were likely due to the fact that with between 1-3x coverage GBS data we were able to determine that two animals are genetically similar, but were not able to obtain a sufficient number of loci with precisely inferred genotyped to find opposing homozygous loci.

Consistent with previous work, we found that using a hand-tuned assignment threshold could reduce the number of false positives (Huisman, 2017; Riester et al., 2009). An alternative approach would be to adaptively determine the assignment threshold via introspection of the underlying data (Grashei et al., 2018). In the majority of the simulations, a fixed threshold of 10 was used based on pilot simulations with array data. As we demonstrate in Figure 5, substantially raising the threshold for assignment could reduce the false-positive rate even for 50,000 markers and low-coverage sequence data, although at the cost of a decreased true-positive rate. The optimal threshold value for assignment depends on the overall sequencing coverage, making it challenging to use a fixed threshold in cases where individuals are sequenced at different coverages. We believe that automating this process is an area for future research, and may depend on the exact breeding program structure, the exact GBS system deployed (e.g., Baird et al., 2008; Davey et al., 2011; Elshire et al., 2011), and reason that parentage information is required.

Furthermore, we believe that the issue of false parent-assignments may be less of an issue in the context of commercial agricultural populations compared to wild populations for two reasons. First, most of the false assignments that we observed were cases where the true parent was not included in the pedigree and a full- or half-sib of the true parent was included and wrongly assigned as a parent. In the context of many animal breeding programs, the routine use of pairs of sibs as parents may not commonly arise because of explicit efforts to manage diversity and inbreeding (e.g., Woolliams et al., 2015). Second, due to the genetic similarity between the full-sib of the true parent and the true parent, using the full-sib of the true parent as a “proxy” parent for the progeny may have limited impact on downstream applications such as estimation of breeding values. Further research is required to quantify the impact of such false positives in downstream applications.

### Constructing proposal distributions via peeling

In this paper, we closely followed the approach of Huisman (2017) for performing parentage assignment, with two differences. First, we modified the genotype probability function to handle sequence data. Second, we recast the construction of proposal distributions for relatives as a series of peeling operations on artificial pedigrees. We believe the later development is of more interest. Peeling provides a rich and computationally efficient framework for estimating the genotypes of a relative based on the genotypes of individuals in an existing pedigree. In this paper we focused on a small number of possible relationships, but this framework can be easily extended to consider a wider and potentially complex class of relatives (e.g., siblings of the focal individual, cousins of the parent, or grandparents), or could be altered to assess alternative relationships (e.g., performing grandparent assignment instead of parentage assignment). Use of these additional relationship classes may depend on the purpose of a particular application.

## Conclusion

In conclusion, we extended the algorithm of Huisman (2017) to perform parentage assignment with sequence data, and evaluated the performance of using low-coverage GBS data for parentage assignment. We found that low-coverage GBS data could be used for accurate parentage assignment, but that there may be concerns with false positives if the true parent is not included on the list of putative parents. Such false positives might be mitigated on a case-by-case basis by tuning the assignment criteria used. These results suggest that GBS data can be used as an alternative to array data for parentage assignment.

## Footnotes

**Conflicts of interest** The authors declare they have no competing interests.

## Acknowledgements

The authors acknowledge the financial support from the BBSRC ISPG to The Roslin Institute BB/J004235/1, from Genus PLC and from Grant Nos. BB/M009254/1, BB/L020726/1, BB/N004736/1, BB/N004728/1, BB/L020467/1, BB/N006178/1 and Medical Research Council (MRC) Grant No. MR/M000370/1. This work has made use of the resources provided by the Edinburgh Compute and Data Facility (ECDF) (http://www.ecdf.ed.ac.uk).

